# Multiscale Modeling of Cancer

**DOI:** 10.1101/033977

**Authors:** Kerri-Ann Norton, Meghan McCabe Pryor, Aleksander S. Popel

## Abstract

Breast cancer remains the second leading cause of cancer death in women, exceeded only by lung cancer. Specifically, triple-negative breast cancer (TNBC) has the worst prognosis, as it is more invasive and lacks estrogen, progesterone, and HER2 receptors that can be targeted with therapies. Due to the need for effective therapies for this type of breast cancer, it is critical to develop methods to (1) understand how TNBC progresses and (2) facilitate development of effective therapies. Here, we describe a multiscale model focusing on tumor formation. Our approach uses multiple scales to investigate the progression and possible treatments of tumors.

## I. Introduction

Cancer is a complex disease that forms and progresses through multiple processes and phases. These processes include sustained tumor growth, metastasis, angiogenesis, and immune response [1]. The complexity of these processes and the involvement of multiple spatial and temporal scales, from the molecular (e.g., genetic and epigenetic alterations, receptor-ligand interactions), to signaling networks, to cellular level (e.g., enhanced proliferation, migration and invasion, epithelial-mesenchymal transition), to tumor/tissue level (e.g., tumor heterogeneity, tumor microenvironment), to whole organism (e.g., circulated tumor cells, mechanistic pharmacokinetics). A number of reviews describe the methodology of multiscale modeling as well as their applications to specific cancer types [2–12]. In this article we present examples of multiscale modeling based on the studies from our laboratory.

One of the proposed mechanisms by which tumors have sustained growth over long periods of time is through cancer stem cells. A subpopulation of cells has the ability to continually divide into two cell types: other stem cells and progenitor cells that have a set number of divisions until they become senescent. This population is then proposed to be the only population that can form metastatic colonies in other organs. An important capability of tumors is their ability to grow initially without a circulatory system that provides oxygenation and a means for cancer cell dissemination. Growth beyond this avascular stage requires tumors to form capillaries from pre-existing vascular structures, that allow for blood flow, in a process known as angiogenesis [4]. A key driving factor in angiogenesis is vascular endothelial growth factor (VEGF). In experimental work done in our laboratory, Lee et al. found that TNBC cell lines secrete cytokine IL6 which primes lymph nodes and lymphatic vessels at metastatic sites (lung) to secrete VEGF [13]. Secretion of this growth factor will in turn recruit vasculature to the metastatic site and prime the site for tumor growth. They also found that the lymphatic vessels primed by IL6 secrete a chemokine CCL5, which recruits the cancer cells to the metastatic site.

Therapies have been developed to target CCL5/CCR5 and VEGF, individually. Maraviroc, which targets CCR5, is an FDA approved drug for HIV. It has yet to be used for cancer but according to our research it shows great promise for a combined therapy. Bevacizumab, an anti-VEGF therapy, initially showed promise for breast cancer but was eventually revoked because it did not improve overall survival. These therapies have been shown to make an improvement in their target areas, however they need to be better understood for use in combination. Our previous experiments have shown that the combination of a VEGF inhibitor with a CCR5 inhibitor was able to substantially reduce the amount of lung metastasis in a spontaneous metastasis xenografts model [13]. Given the poor prognosis of TNBC and urgent demand for effective therapies, there is a need to be able to assemble our current understandings of tumor growth, along with angiogenesis, in an efficient manner. One way to tackle this task is by building a computational systems biology model.

Multiscale computational models are an important tool that can be used to study complex mechanisms on a variety of scales and deduce how each level impacts the other. Multiscale models can span two or more scales, depending on the problem at hand. Below we describe a multiscale agent-based model that investigates TNBC tumor growth based on CCR5 presence and how inhibition of CCR5 impacts that tumor growth. This is an extension of the model [14]. We then discuss further extension of this model to include vascularization based on VEGF concentrations and prediction of how anti-angiogenic therapies would impact tumor growth.

## II. Results

### A. Avascular Tumor Growth

The agent-based model of avascular tumor growth was developed to simulate a metastatic TNBC cell line (MDA-MB-231 abbreviated here as MB231) and a more metastatic cell line that had been propagated to metastasize to the lung (MB231-luc). The model spans multiple scales by using the molecular (CCR5) and cellular (stem and progenitor) scales in combination to predict the growth of the tissue scale (Tumor). Overall, we found that the growth rate of the MB231-luc tumor was greater than the MB231 tumor. We also determined that the growth of the overall tumor, both in rate and in distribution, was correlated with the growth and distribution of the CCR5+ cells. Finally, we observed that the simulated tumor had a finger-like architecture, as seen in Fig. 1, which is frequently seen in invasive tumors.

**Figure 1:**
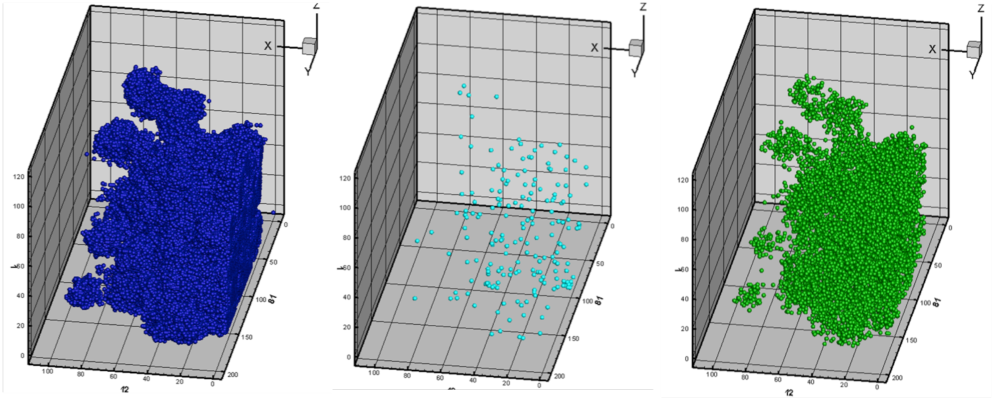
Simulation of MB231-luc tumor growth. The cells in blue are the total tumor, the cyan cells are the stem cells and the green cells are the CCR5+ cells. The growth shows the finger-like morphology characteristic of invasive growth.

### B. CCR5 Inhibitor Therapy

In addition to the tumor growth model, we ran simulations that included the impacts of Maraviroc, a CCR5 inhibitor. We found that the addition of Maraviroc did not slow the growth rate of the tumor, similar to in vivo experiments [13]. We speculate that while this treatment does not affect the primary growth of the tumor, it would lower the rate of invasion and metastasis.

### C. Tumor Angiogenesis and Anti-Angiogenic Therapies

While the current model has shown us the impact of CCR5 on tumor growth, an interesting extension would be to include the influence of VEGF and angiogenesis on tumor growth, as well. Our previous experimental work has indicated that TNBC cells are able to stimulate VEGF secretion at the metastatic site, which in turn stimulates angiogenesis. Inclusion of VEGF dependent angiogenesis in this model would allow us to gain understanding into how vascularization impacts tumor growth. The model would also give insights about the effect of anti-angiogenic therapies and combination therapies of CCR5 inhibitors with anti-angiogenic drugs.

## III. Methods

### A. Decision Tree

Agent-based models represent cancer cells as agents that make decisions of which cellular functions (proliferation, migration, senescence, etc.) to do with the aid of a decision tree. These decisions are based on their cellular properties, environment, and probabilities as seen in Fig. 2

**Figure 2:**
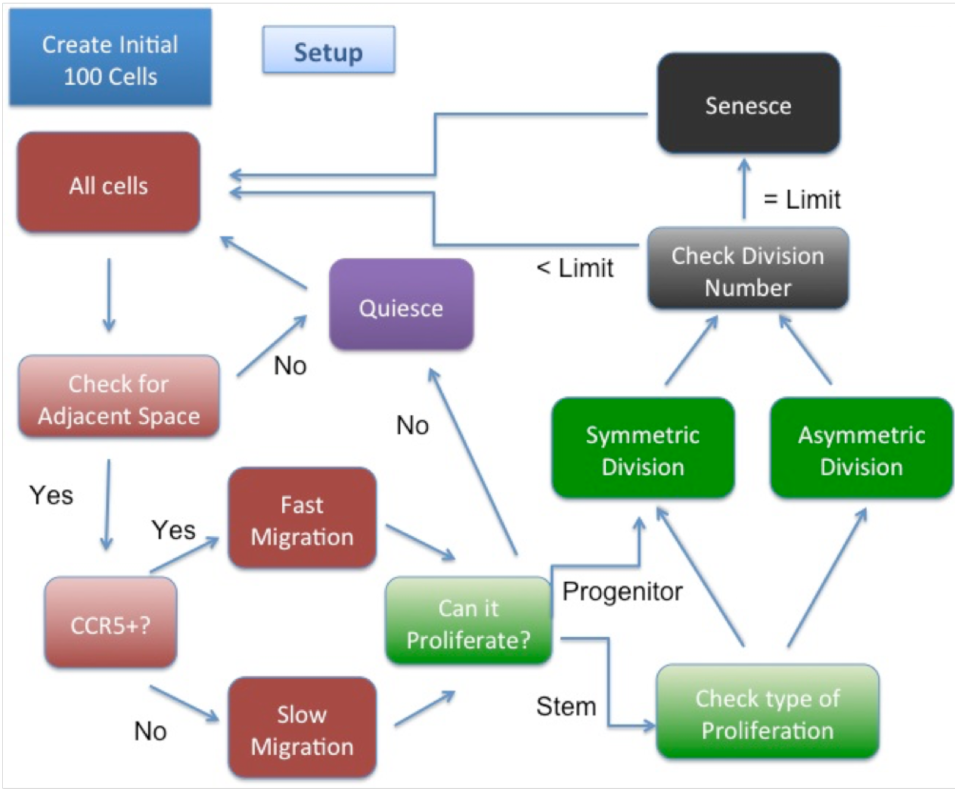
Decision tree for tumor model.

In the model we assume that stem cells can reproduce indefinitely to produce either another stem cell or a progenitor cell. Progenitor cells can only proliferate 12 times before they become senescent and die. CCR5+ cells migrate 4 times as fast as CCR5-cells. If a cell is surrounded by other cells it becomes quiescent and can no longer proliferate or migrate until there is free space. We allow the simulation to run for 3 years of simulation time or until the tumor reaches 500,000 cells.

### B. Simulation Space and Initial Conditions

We used flow cytometry in two different triple-negative cell lines using MDA-MB-231, a primary cell line, and MDA-MB-231-luc, a cell line expressing luciferase and selected to metastasize to the lymph nodes to determine the percentages of different cell populations. In particular, we measured cell surface receptors CCR5, CD24, and CD44 to determine the numbers of CCR5 high cells and the number of stem cells in each cell line. These measurements were then input into the model.

The model starts with a distribution of 100 cells each with stem cells and CCR5 cells at different percentages for each cell line. 100 cells are placed in the gridspace (500*500*500) in the proportions that are found in each cells line. Specifically in the MB231 case, there are 90 CCR5-progenitor cells, 8 CCR5-stem cells, 1 CCR5+ cell and 1 CCR5+ stem cell. In the MB231-luc case there are 75 CCR5-progenitor cells, 19 CCR5-stem cells, 5 CCR5+ cells and 1 CCR5+ stem cell.

### C. Model Assumptions

There are many assumptions made to the model to simplify the complexity of cancer. For one, we assume that this is an avascular tumor, before the initiation of angiogenesis and thus we assume that all the cells have enough oxygen to survive and proliferate. This is why the model is stopped at 500,000 cells which is the roughly the size it would start the initiation of angiogenesis. We also assume that space limitation is not caused by the organ itself but only the cancer cells; this is a realistic assumption because tumors tend to be much denser than the surrounding breast tissue. Another assumption we make is that at this stage in the growth the CCR5+ cells have an increase in random motility not chemotaxis, once again due to the fact that the tumor is small and has not secreted enough factors to trigger the lymph nodes to secrete CCL5; CCL5 gradients should be included in future models.

### D. Areas Where Agent-Based Modeling is Useful

Agent-based models are extremely useful when studying the dynamics of tumor growth. Due to the probabilistic nature of this model type, agent-based models are very versatile to the system and scale that can be considered. This style model is also useful in many other areas including other forms of cancer beyond TNBC, modeling epidemic disease spread, and immune cell dynamics. The power of agent-based models lies in the fact that one can use a simplified version of cancer to form hypotheses as to what mechanisms are essential for tumor growth patterns that are seen and which mechanisms are only peripheral to the system being studied. Using these types of models we can also predict the most effective therapy combinations and treatment schedules that can then be tested in vivo.

